# Systematic discovery of plasmid-dependent inoviruses highlights broad infectivity across divergent IncF plasmids

**DOI:** 10.64898/2026.11.23.768622

**Authors:** Eva Githuku, Hannah R. Kumar, Rhiannon T. Mariano, Alecia B. Rokes, Erica Lasek-Nesselquist, Siân V. Owen

## Abstract

Plasmid-dependent phages (PDPs) exploit bacterial conjugation machinery to infect their hosts, yet their natural diversity and specificity for different conjugative plasmids remain poorly understood. Here, we extend Phage DisCo, a fluorescence-based differential plating assay, to enable the systematic and selective isolation of distinct groups of F-dependent phages. By combining RNase treatment with deletion of the bacterial *tolA* gene, we established screening conditions that distinguish single-stranded RNA phages, filamentous single-stranded DNA inoviruses, and phages with alternative infection requirements. Applying this approach to environmental samples, we established a collection of 27 plasmid-dependent inoviruses isolated using two divergent IncF conjugative plasmids, F and pED208, whose conjugation proteins share only 30-50% amino acid sequence identity. Strikingly, all 27 isolates infected bacteria carrying either plasmid, regardless of the plasmid used for their initial isolation. Whole-genome phylogenetic analysis revealed that phages recovered using F and pED208 were interspersed throughout the phylogeny rather than forming distinct plasmid-associated lineages. These findings demonstrate that naturally occurring F-dependent inoviruses comprise a diverse population capable of exploiting divergent conjugation systems, despite differences in their component proteins. Our results establish a selective approach for investigating plasmid-dependent phage diversity and highlight the importance of systematic culture-based isolation for understanding the diversity and host specificity of plasmid-dependent phages.

## INTRODUCTION

Conjugative plasmids are major drivers of horizontal gene transfer in bacteria, facilitating the dissemination of antibiotic resistance genes and other adaptive traits^1^. These plasmids encode type IV secretion systems (T4SSs) that mediate plasmid DNA transfer between bacterial cells, producing extracellular pili that facilitate contact between donor and recipient cells^2^. Conjugative pili can also serve as receptors for bacteriophages known as plasmid-dependent phages (PDPs)^3,4^. The F plasmid of *Escherichia coli* is among the best-characterized examples, with several evolutionarily distinct groups of phages known to exploit its conjugation pilus as a receptor. These include single-stranded RNA (ssRNA) phages such as MS2 and Qβ (family *Fiersviridae*), filamentous single-stranded DNA (ssDNA) phages such as f1, fd, and M13 (family *Inoviridae*), and the rare, more recently described tailed double-stranded DNA (dsDNA) phages FtMidnight and PDP46^4,5^. ssRNA F-plasmid dependent phages are collectively termed Fr phages, whilst ssDNA filamentous F-plasmid dependent phages are known as Ff phages. Although these phages share a dependence on conjugative plasmids, they represent completely distinct evolutionary groups, and differ in their infection mechanisms and requirements for additional host factors^6,7^.

The F-dependent Inoviridae were among the first plasmid-dependent phages discovered^8–10^, and the closely related phages f1, fd, and M13 have since become extensively studied as models of bacteriophage biology and are the foundation of phage display technology^11,12^. However, these model phages represent only a small fraction of the natural diversity of inoviruses. A large-scale genomic and metagenomic survey identified more than 10,000 putative inovirus sequences, revealing a vast diversity that remains largely unexplored experimentally^13^. The extent to which this genomic diversity is reflected among infectious plasmid-dependent inoviruses, and the range of conjugative plasmids they can exploit, remain poorly understood. Conjugative plasmids encode related but often highly divergent conjugation systems^14–16^, raising the question of whether this divergence restricts the host range of the phages that depend on them.

We previously developed Phage DisCo, a fluorescence-based differential plating assay for the systematic isolation of plasmid-dependent phages from environmental samples^4^. Here, we extend Phage DisCo to enable the systematic and selective isolation of different groups of F-dependent phages. We isolated 27 inoviruses using the divergent F-like conjugative plasmids F and pED208. All isolates were capable of infecting hosts carrying either plasmid, and comparative genomic analysis revealed a shared population of closely related inoviruses rather than distinct plasmid-associated lineages. These findings demonstrate the utility of selective culture-based discovery for investigating plasmid-dependent phage diversity and specificity.

## RESULTS

### Selective screening distinguishes major groups of F-dependent phages

In its original implementation, Phage DisCo used two taxonomically distinct bacterial hosts carrying the same conjugative plasmid, each labeled with a different fluorescent protein, to distinguish plasmid-dependent phages (PDPs) from plasmid-independent phages^4^. Here, we modified this approach to use two otherwise isogenic bacterial backgrounds that differ in conjugative plasmid carriage, each labeled with a distinct fluorescent protein^17^. This eliminates the requirement for phages to infect two distinct bacterial backgrounds, enabling the detection of PDPs with narrower bacterial host ranges. When combined in a mixed bacterial lawn, PDPs selectively infect the plasmid-bearing strain, producing plaques with a single fluorescent signal, whereas plasmid-independent phages capable of infecting both strains produce plaques lacking fluorescence. This modified plasmid-dependent phage DisCo approach is similar to an approach recently used to identify novel plasmid targeting alphatectiviruses^18,19^.

To enable the selective isolation of different groups of F-dependent phages, we exploited differences in their infection requirements. Supplementation of the plating medium with RNase was used to suppress ssRNA phages^20^, while deletion of the host gene *tolA*, which is required for infection by F-dependent inoviruses^7,21^, was used to exclude filamentous ssDNA phages (Fig. 1a). Combining these approaches should enable the detection of rare F-dependent phages that might otherwise be obscured by more abundant ssRNA and filamentous ssDNA phages in conventional Phage DisCo assays, including tailed dsDNA phages such as FtMidnight.

**Figure 1.**
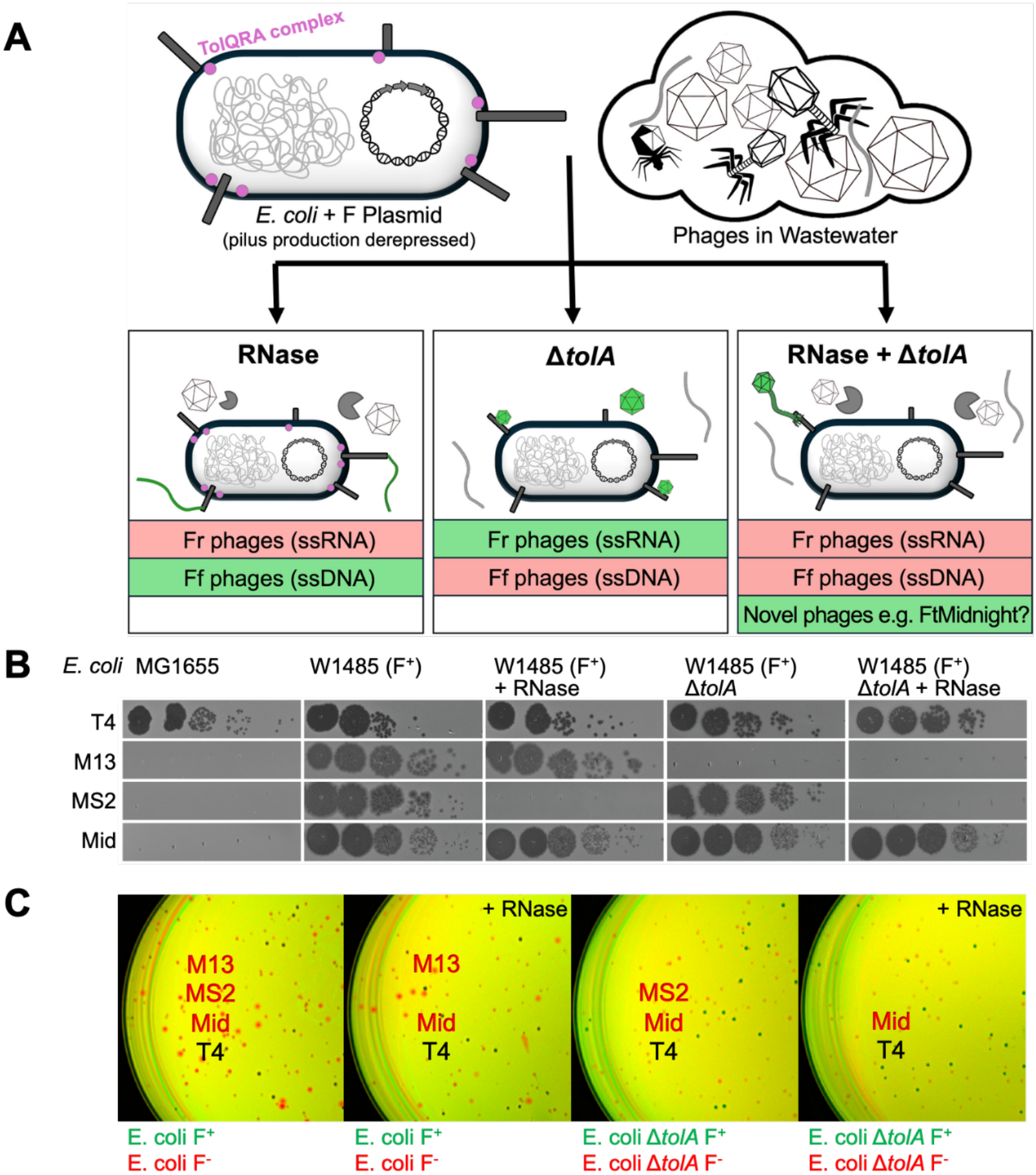
Selective detection of plasmid-dependent phages using the Discovery by Coculture (DisCo) assay. **(A)** Schematic illustrating the sensitivity of representative bacteriophages to RNase treatment and deletion of the host *tolA* gene. RNase treatment inhibits ssRNA phages (MS2), whereas deletion of *tolA* prevents infection by inoviruses (M13). Combining both conditions enables the detection of phages that are neither RNase-sensitive nor TolA-dependent, represented by FtMidnight. Green and red boxes indicate phage groups expected to be detected or inhibited, respectively, under each screening condition. The somatic phage T4 is unaffected by either treatment. **(B)** Representative plaque assays demonstrating the susceptibility of M13, MS2, FtMidnight, and T4 to RNase treatment, deletion of *tolA*, or both conditions combined. **(C)** Representative DisCo assays using the four model phages under the different selective conditions. Fluorescently labeled F^+^ and F^−^ *E. coli* strains are combined to distinguish plasmid-dependent phages (red plaques) from plasmid-independent phages (black plaques). RNase treatment and deletion of *tolA*, individually or in combination, selectively eliminate plaques formed by different classes of PDPs while retaining detection of phages that are insensitive to both conditions.

We evaluated the selectivity of these screening conditions using three representative F-dependent phages with distinct genome types: MS2 (ssRNA), M13 (ssDNA), and FtMidnight (dsDNA), alongside the plasmid-independent phage T4 (Fig. 1b,c). As expected, supplementation with RNase prevented plaque formation by MS2 but not M13 or FtMidnight, whereas deletion of *tolA* prevented infection by M13 but not MS2 or FtMidnight. Combining RNase treatment with Δ*tolA* prevented plaque formation by both MS2 and M13 while retaining susceptibility to FtMidnight. Infection by the plasmid-independent phage T4 was unaffected by RNase treatment and *tolA* deletion. Together, these results demonstrate that RNase treatment and *tolA* deletion can be used individually or in combination to selectively recover different groups of F-dependent phages.

We next applied the four screening conditions to wastewater samples to investigate the relative recovery of different groups of F-dependent phages (Supplementary Fig. 1). Although total plaque numbers varied between samples, RNase treatment generally resulted in a greater reduction in PDP plaques than deletion of *tolA*, suggesting that ssRNA phages accounted for a larger proportion of recoverable F-dependent phages than inoviruses in these samples. Notably, no PDP plaques were detected under the combined RNase and Δ*tolA* conditions in any of the samples tested in this study, suggesting that F-dependent phages other than ssRNA phages and TolA-dependent inoviruses were rare or absent from these samples.

### Plasmid-dependent inoviruses infect hosts carrying divergent F-like conjugative plasmids

The ability to selectively isolate different groups of plasmid-dependent phages (PDPs) provides an opportunity to systematically investigate their natural diversity and host specificity. We chose to focus on inoviruses, using RNase-supplemented Phage DisCo to investigate their diversity and host range across two divergent F-like conjugative plasmids: the classical F plasmid and pED208, a derepressed derivative of the naturally occurring F-like plasmid F_0_lac^15^. Despite encoding functionally related conjugation systems, homologous Tra/Trb proteins from F and pED208 generally share only 30–50% amino acid sequence identity (Fig. 2a). Notably, TraA, the major structural subunit of the conjugative pilus, shares only 32.3% amino acid sequence identity between the two plasmids. Screening of environmental samples yielded numerous plasmid-dependent plaques, from which we selected 27 phages for isolation and further characterization, including 21 recovered using F and six using pED208 (Fig. 2b, Supplementary Tables S1 and S2).

**Figure 2.**
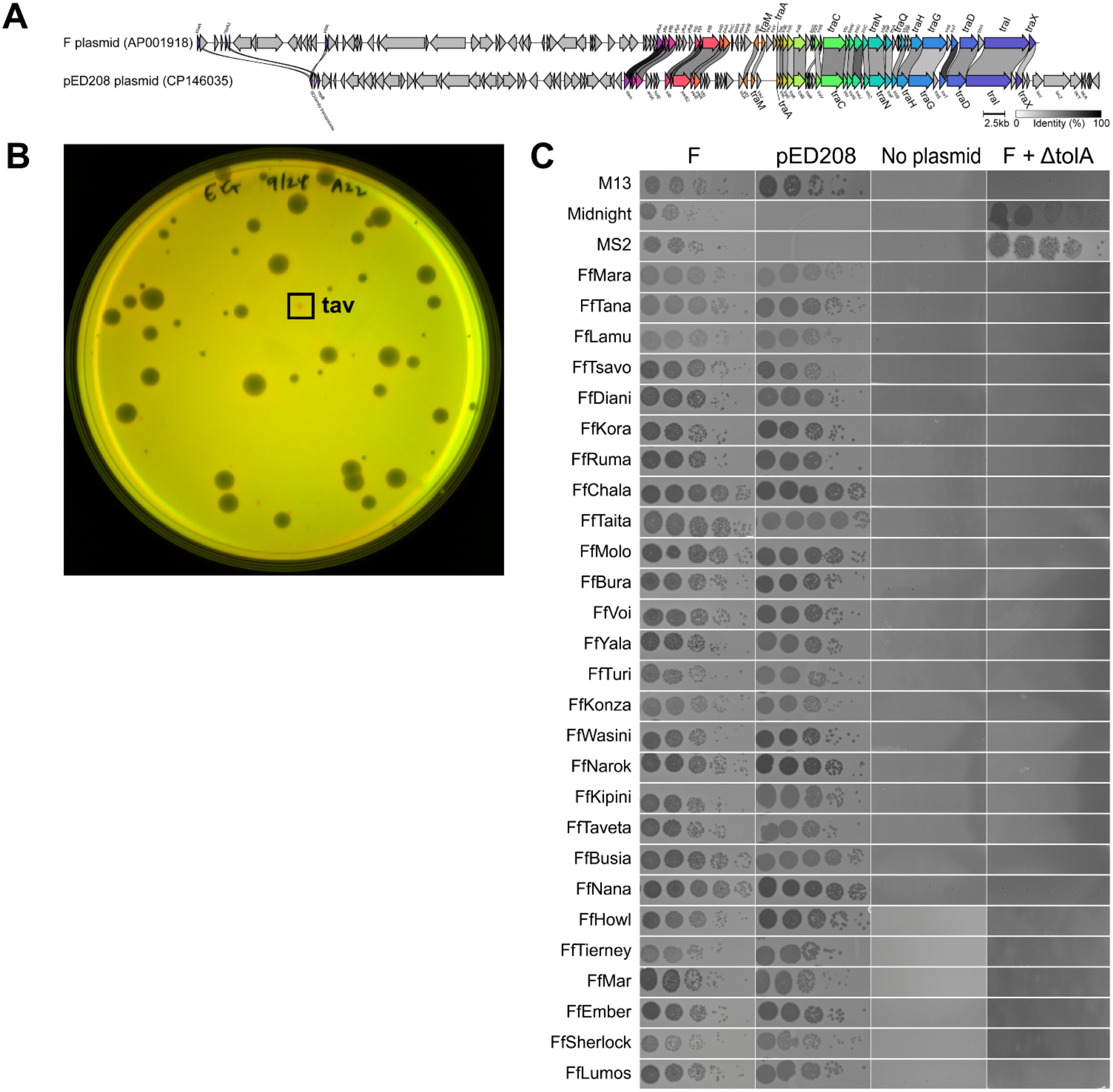
Isolation and characterization of plasmid-dependent inoviruses using the Discovery by Coculture (DisCo) assay. (A) Comparison of the genetic organization of the F and pED208 conjugative plasmids, with amino acid sequence identities indicated for homologous proteins. (B) Representative DisCo assay showing the detection of plasmid-dependent phages as red plaques. The boxed plaque was picked for isolation and subsequently designated FfTaveta. (C) Serial-dilution spot assays validating the infection phenotypes observed during DisCo isolation for all isolated inoviruses and three reference phages. Phages were tested against four *E. coli* host backgrounds: W1485 carrying the F plasmid, BW25113 carrying pED208, plasmid-free MG1655, and W1485 Δ*tolA* carrying the F plasmid. These assays confirm plasmid-dependent infection, cross-infectivity between F- and pED208-containing hosts, and dependence on host TolA.

To determine whether these phages were specific to the plasmid used for their isolation, we assessed their ability to infect *E. coli* carrying F, pED208, or no conjugative plasmid. Strikingly, all 27 isolates infected hosts carrying either F or pED208, regardless of the plasmid used for their initial isolation, whereas none infected plasmid-free cells (Fig. 2c). Furthermore, none of the isolates infected the corresponding Δ*tolA* hosts, consistent with the TolA dependence characteristic of F-dependent inoviruses. Together, these findings demonstrate that naturally occurring plasmid-dependent inoviruses can infect hosts carrying divergent F-like conjugative plasmids, revealing a lack of strict plasmid specificity despite substantial divergence in their conjugation machinery.

### Genomic diversity of plasmid-dependent inoviruses

To investigate the evolutionary relationships among the isolated inoviruses, we sequenced all 27 isolates and compared their genomes with those of previously characterized F-dependent inoviruses. All 27 isolates possessed closely related genomes ranging from 6,393 to 6,412 bp in length, with pairwise nucleotide sequence identities ranging from 89.5% to greater than 99.9%, and shared the same protein-coding gene repertoire and gene synteny as the well-characterized filamentous phage M13. We assigned the isolates the prefix Ff (F-dependent filamentous phage), consistent with the established designation for filamentous phages that infect F-plasmid-bearing bacteria^22^.

Whole-genome phylogenetic analysis resolved three major clades, with phages isolated using F and pED208 interspersed throughout the phylogeny rather than forming distinct plasmid-associated lineages (Fig. 3). Together with their shared ability to infect hosts carrying either plasmid, these findings suggest that the isolates represent a common population of inoviruses capable of exploiting divergent F-like conjugation systems, rather than distinct populations specialized for individual plasmids.

**Figure 3.**
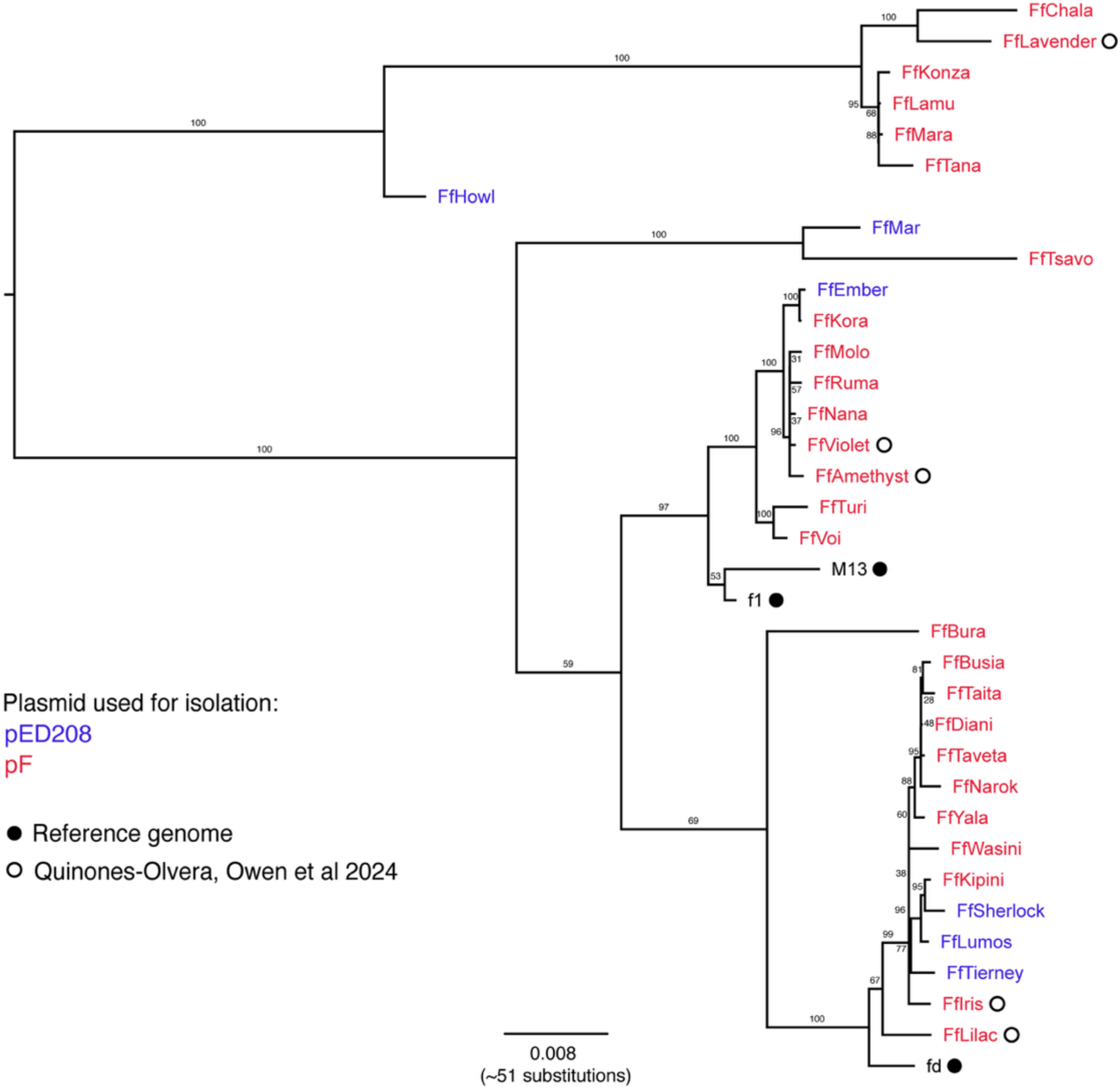
Phylogenetic relationships among plasmid-dependent inoviruses isolated using F and pED208 conjugative plasmids. Maximum-likelihood phylogenetic tree based on whole-genome nucleotide sequences of inoviruses isolated using hosts carrying the F plasmid (red) or pED208 (blue), alongside reference inovirus genomes. The tree was midpoint-rooted. Phages isolated using the two plasmid backgrounds are interspersed throughout the phylogeny, indicating that F and pED208 support infection by overlapping populations of inoviruses.

### Evolutionary relationships among individual inovirus proteins

To investigate whether individual phage proteins shared the evolutionary relationships observed at the whole-genome level, we constructed phylogenetic trees for seven conserved proteins and compared their topologies with the whole-genome phylogeny (Fig. 4). Phylogenies of the replication (Fig. 4A), assembly, attachment, major coat, and minor coat proteins broadly recapitulated the three clades identified in the whole-genome analysis (Supplementary Fig. 3). In contrast, the single-stranded DNA-binding (Fig. 4B) and morphogenesis proteins exhibited greater differences in tree topology, with isolates grouping differently from their positions in the whole-genome phylogeny. These findings suggest that evolutionary relationships are largely conserved across the inovirus proteome, although discordance among individual protein phylogenies points to potentially distinct evolutionary histories for some proteins.

**Figure 4.**
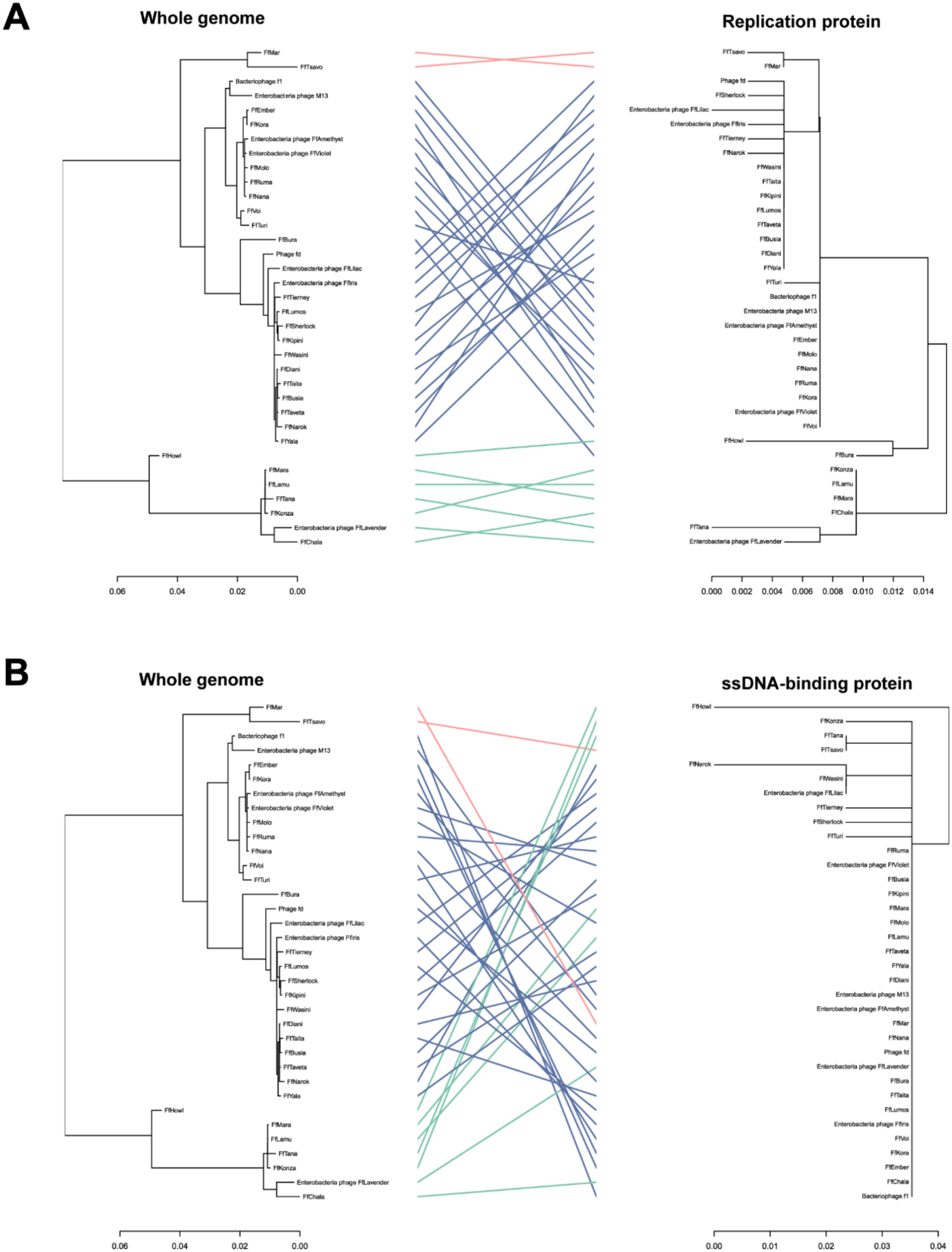
Comparison of whole-genome and individual protein phylogenies of plasmid-dependent inoviruses. Tanglegrams comparing phylogenetic relationships inferred from complete inovirus genomes with those inferred from individual phage proteins: (A) replication protein and (B) single-stranded DNA-binding protein. In each panel, the whole-genome phylogeny is shown on the left and the corresponding protein phylogeny on the right. Lines connect matching phages between trees and are colored according to the three major clades identified in the whole-genome phylogeny. Crossing lines indicate differences in phylogenetic topology between the whole-genome and individual protein trees.

## DISCUSSION

Plasmid-dependent phages exploit bacterial conjugation machinery to infect their hosts, yet the natural diversity and host specificity of these phages remain poorly understood. Here, we extended Phage DisCo to enable the systematic and selective isolation of different groups of F-dependent phages, using differences in their infection requirements to distinguish ssRNA phages, TolA-dependent inoviruses, and phages with alternative infection requirements.. Applying this approach to environmental samples, we recovered 27 plasmid-dependent inoviruses and investigated their genomic diversity and ability to infect bacteria carrying divergent F-like conjugative plasmids. Despite substantial sequence divergence between the conjugation systems of F and pED208, all 27 isolates infected hosts carrying either plasmid. Moreover, isolates recovered using the two plasmids were interspersed throughout the phylogeny, indicating that they belong to an overlapping population rather than any distinct plasmid-associated lineages.

The selective screening approach developed here provides a means of investigating groups of plasmid-dependent phages (PDPs) that might otherwise be obscured by more abundant or readily isolated phages. In our wastewater samples, RNase treatment generally reduced the number of recoverable F-dependent phages more substantially than deletion of *tolA*, suggesting that ssRNA phages were more prevalent than inoviruses under our screening conditions. Notably, no PDPs were detected in any of the 108 wastewater samples screened using the combined RNase and Δ*tolA* conditions, despite our demonstration that this approach permits infection by the recently discovered tailed F-dependent phage FtMidnight. These findings suggest that non-canonical F-dependent phages, including FtMidnight-like phages and potentially other previously uncharacterized groups, are extremely rare or absent from the wastewater samples examined here. Interestingly, FtMidnight and PDP46 were originally isolated from farm compost^4^ and shellfish^5^, respectively, raising the possibility that these phages are more readily recovered from environments other than wastewater. Expanding selective screening to a wider range of environmental sources may therefore be particularly valuable for uncovering additional diversity beyond the well-characterized ssRNA and filamentous ssDNA F-dependent phages. As our understanding of the molecular requirements for PDP infection continues to expand^23^, this knowledge can be leveraged to design increasingly selective Phage DisCo screens, enabling the systematic isolation and characterization of previously unexplored groups of plasmid-dependent phages.

A striking finding of this study was the ability of all 27 isolated inoviruses to infect bacteria carrying either F or pED208, despite the substantial divergence between their conjugation systems. Although the ability of filamentous F-dependent phages to infect hosts carrying different F-like plasmids has been recognized previously^24,25^, our systematic isolation of naturally occurring inoviruses demonstrates that this broad infectivity is shared across a much larger collection of phages. The absence of plasmid-associated clustering in the whole-genome phylogeny further supports the conclusion that the two plasmids are susceptible to a common population of inoviruses, rather than distinct populations specialized for individual conjugation systems. These observations suggest that the sequence divergence between F-like conjugation machineries does not necessarily translate into corresponding barriers to infection by plasmid-dependent inoviruses.

The molecular basis of this broad infectivity remains unclear. F-dependent inoviruses initiate infection through interactions with the conjugative pilus, followed by engagement of the host TolQRA complex to facilitate entry across the bacterial envelope^7^. Recent structural studies of conjugative pilus tips have revealed a complex architecture involving the accessory proteins TraG and TraH, in addition to the major pilin subunit^26,27^. These findings raise the possibility that inoviruses recognize conserved structural features of the pilus tip, rather than the highly divergent pilin subunits themselves. However, TraG and TraH also exhibit substantial sequence divergence between F and pED208, suggesting that any conserved recognition features may be maintained at the structural rather than primary sequence level. Alternatively, inoviruses may tolerate substantial variation in their receptor interactions. Identifying the molecular determinants of pilus recognition will be important for understanding how these phages infect hosts carrying divergent conjugative plasmids.

Whole-genome phylogenetic analysis resolved the isolated inoviruses into three major clades, despite their shared ability to infect both F- and pED208-bearing hosts. Most individual protein phylogenies broadly recapitulated these relationships, suggesting that evolutionary patterns are largely conserved across the genome. Thus, the genetic variation observed among these isolates does not appear to be associated with differences in their qualitative plasmid host range, although differences in infection efficiency may exist.

Together, our findings demonstrate the utility of selective Phage DisCo screening for systematically investigating plasmid-dependent phage diversity and reveal broad infectivity across divergent F-like conjugative plasmids among naturally occurring inoviruses. The substantial differences between the inovirus diversity revealed by metaviromic surveys^13^ and that represented by experimentally isolated phage genomes highlight the limitations of existing culture-based collections. Developing systematic isolation approaches that capture a broader range of viral diversity will be important for bridging this gap more broadly and enabling experimental investigation of phages currently identified only from sequence data.

## METHODS

### Bacterial strains and growth conditions

Bacterial strains and plasmids used in this study are listed in Supplementary Tables S3 and S4. Unless otherwise indicated, *Escherichia coli* strains were grown in LB Lennox broth with aeration (200 rpm) or on LB agar plates at 30°C or 37°C, as appropriate. Where required, antibiotics were added at the following final concentrations: chloramphenicol (20 µg/mL), kanamycin (50 µg/mL), and tetracycline (20 µg/mL).

### Phage propagation and titration

Bacteriophages used in this study and their corresponding propagation hosts are listed in Supplementary Table S1. High-titer phage stocks were prepared by inoculating exponentially growing host cultures (approximately OD_600_ = 0.3) with approximately 10^6^ plaque-forming units (PFU) of phage. Cultures were incubated at 37°C with aeration overnight, after which bacterial cells were removed by centrifugation at 12,000 g for 2 minutes. Supernatants were passed through 0.22-µm filters to obtain cell-free phage stocks.

Phage titers were determined using double-layer agar spot-titer assays. Briefly, 100 µL stationary-phase host cultures were mixed with 3 mL molten LB top agar (0.5% agar) and poured onto LB agar plates. Phage stocks were serially diluted tenfold in LB broth, and 3 µL of each dilution was spotted onto the solidified bacterial lawns. Plates were incubated at 37°C overnight before plaques were enumerated.

### Strain construction

Fluorescent *E. coli* strains were generated by introducing plasmids encoding constitutively expressed sGFP2 or mScarlet-I, together with a chloramphenicol resistance marker, by electroporation.

Chromosomal deletions of *tolA* were generated using λ-Red recombineering. Briefly, the plasmid pSIM5-tet was introduced into *E. coli* by electroporation and growth at 30°C, and expression of the λ-Red recombination proteins was induced by incubation at 42°C for 15 min. Cultures were subsequently chilled on ice and prepared for electroporation. Approximately 100 ng of purified PCR product was introduced into 50 µL of electrocompetent cells via electroporation. Following recovery, transformants were plated on selective LB agar at 37°C to isolate recombinant colonies that had lost the pSIM5-tet plasmid.

Conjugative plasmids were transferred between *E. coli* strains by solid-surface mating. Briefly, donor and recipient strains were grown overnight in LB Lennox broth at 37°C with appropriate antibiotic selection. Equal volumes of donor and recipient cultures were mixed, pelleted by centrifugation at 3,000 × *g* for 5 min, and resuspended in a small volume of LB broth. Cell mixtures were spotted onto LB agar and incubated at 37°C for 4 h to facilitate conjugation. Cells were subsequently recovered and plated to identify transconjugants. For the F plasmid, a derivative carrying a kanamycin resistance marker was used, allowing transconjugants to be selected on kanamycin-containing agar. As pED208 does not encode an antibiotic resistance marker, but encodes a *lac* operon, conjugation mixtures were plated on LB agar supplemented with X-gal and IPTG, and dark blue colonies were picked as putative transconjugants. Acquisition of pED208 was subsequently confirmed by susceptibility to the plasmid-dependent inovirus M13.

### Validation of selective Phage DisCo screening

The selectivity of Phage DisCo screening conditions was evaluated using three F-dependent phages, MS2, M13, and FtMidnight, and the plasmid-independent phage T4. Mixed bacterial lawns were prepared by combining 100 µL each of overnight cultures of *E. coli* MG1655 F912 pEB2-cmR-sGFP and MG1655 pEB2-cmR-mScarlet-I with 3 mL of molten LB top agar. 10 µL of each diluted phage stock (containing approximately 50 PFU) was added to the mixture before pouring onto LB agar plates. The assay was repeated with the addition of 5 µL of RNase A (NEB T3018L, 0.1 mg total used per plate), using the corresponding Δ*tolA* strains, or with both modifications. Plates were incubated overnight at 37°C and imaged using a Bio-Rad ChemiDoc fluorescence imager.

To assess phage susceptibility across the four screening conditions, MS2, M13, FtMidnight, and T4 were serially diluted tenfold in LB Lennox broth. Mixed bacterial lawns were prepared as described above, without the addition of phage. Following solidification, 3 µL of each phage dilution was spotted onto the lawns. Plates were incubated overnight at 37°C and imaged using the Bio-Rad ChemiDoc MP fluorescence imager (fluorescent images) or the Epson Perfection V850 Pro Scanner.

### Collection and processing of environmental samples

Wastewater samples were obtained from the Wastewater Laboratory at the Wadsworth Center (Albany, NY, USA), as part of the New York State wastewater surveillance program. Sampling locations are listed in Supplementary Table S2. Wastewater samples were passed through a 0.22-µm filter to remove bacterial cells. Filtered samples were stored at 4°C until phage screening. All samples were screened within 2 months of collection.

### Isolation of plasmid-dependent phages by Phage DisCo

Plasmid-dependent phages were isolated from filtered wastewater samples using Phage DisCo, as described above. Mixed bacterial lawns were prepared using fluorescently labeled *E. coli* MG1655 strains carrying either the F plasmid (F912) or no conjugative plasmid. For each assay, 100 µL of filtered wastewater was added to the bacterial suspension before pouring the top agar onto LB agar plates. RNase was included in the top agar, as described above, to selectively isolate inoviruses by suppressing ssRNA phages. The same screening procedure was performed using the F-like plasmid pED208 and its corresponding plasmid-free control strain. Phage plaques in individual wastewater samples were manually counted, analyzed in R^28^ (v4.5.1), and visualized with ggplot2^29^ (v4.0.2).

Putative plasmid-dependent phages were identified by plaques exhibiting red fluorescence and isolated by plaque picking. Individual isolates were purified through three successive rounds of plaque isolation. To confirm their infection requirements, putative plasmid-dependent phages were additionally screened in the presence or absence of RNase and using the corresponding Δ*tolA* strains.

Phages isolated by EG were named after locations in Kenya, while the six phages isolated by RM were named independently.

### Phage DNA extraction and sequencing

Inovirus replicative-form DNA was extracted from infected *E. coli* cultures. Briefly, 2 µL of an overnight host culture and 20 µL of phage stock was inoculated into 2 mL of LB Lennox broth and incubated at 37°C overnight. To minimize plasmid DNA contamination during extraction, host strains lacking the small plasmids encoding fluorescent reporters were used. Cells were collected by centrifugation at maximum speed for 1 min, and double-stranded replicative-form DNA was extracted from the cell pellet using a plasmid miniprep kit (New England Biolabs).

DNA concentration and purity were assessed using a NanoDrop spectrophotometer and Qubit dsDNA Broad Range (BR) Assay Kit, and DNA fragment sizes were estimated by agarose gel electrophoresis. Extracted DNA was submitted to Plasmidsaurus for long-read sequencing and assembly using Oxford Nanopore Technologies (Standard Low Concentration Plasmid sequencing service).

### Inovirus genome assembly and phylogenetic analysis

Inovirus genome assemblies were provided by Plasmidsaurus. The completeness and potential contamination of assembled inovirus genomes were assessed using CheckV v1.0.3^30^. Genomes recovered in this study and reference inovirus genomes obtained from NCBI were annotated using Pharokka v1.9.1^31^. Specifically, coding sequences (CDS) were predicted with PHANOTATE^32^. Functional annotation was generated by matching each CDS to the PHROGs database^33^ using MMseqs2^34^ and PyHMMER^35^. Genome sequences were reoriented using Dnaapler^36^ v 1.4.0, with the replication-associated protein gene used as the reference for genome orientation.

Whole-genome sequences were aligned using MAFFT^37^ v7.453, and the resulting multiple sequence alignment was manually inspected using SeaView v5.1^38^. Maximum-likelihood phylogenetic trees were reconstructed using IQ-TREE^39^ v1.6.12 with 1,000 bootstrap replicates and most appropriate substitution model identified by ModelFinder^40^. Trees were visualized in FigTree v1.4^41^.

To investigate evolutionary relationships among individual inovirus proteins, phylogenetic trees were also reconstructed from individual protein sequences. Congruence between the whole-genome and individual protein phylogenies was assessed using the R^28^ (v4.5.1) package dendextend^42^ v1.19.1. The EMBOSS NEEDLE tool was used to calculate pairwise protein identities.

## SUPPLEMENTARY MATERIAL

**Supplementary Figure S1**. Recovery of plasmid-dependent and somatic phages from wastewater samples under different selective screening conditions.

**Supplementary Figure S2**. Original isolation plaques of the 27 plasmid-dependent inoviruses recovered in this study.

**Supplementary Figure S3**. Comparison of whole-genome and individual protein phylogenies of plasmid-dependent inoviruses. Tanglegrams comparing the whole-genome phylogeny with phylogenies reconstructed from individual inovirus protein sequences.

**Supplementary Table S1**. Bacteriophages and corresponding genome sequences used in this study.

**Supplementary Table S2**. Wastewater samples used in this study.

**Supplementary Table S3**. Bacterial strains used in this study.

**Supplementary Table S4**. Plasmids used for the construction of mutant strains in this study.

**Supplementary Table S5**. Primers used in this study

## ACKNOWLEDGEMENTS

We are grateful to Jonathan Plitnick, Monica Foote and all staff at the Wastewater laboratory in the Wadsworth Center for assistance with wastewater samples. We also thank the staff at the Wadsworth Center Glassware and Media cores for technical support. We thank Jonasz Patkowski and Thiago Costa for providing the pED208 plasmid. This study was supported by the Centers for Disease Control and Prevention (CDC) Epidemiology and Laboratory Capacity (ELC) Program, funded through Health Research, Inc (HRI) and the New York State Department of Health (NYSDOH) with Federal Award Identification Number NU50CK000516. We thank the WWTP operators who pull samples for the New York state wastewater surveillance network. The contents are those of the author(s) and do not necessarily represent the official views of, nor an endorsement by, NYSDOH, HRI, or CDC.

HRK and ABR were supported by the Public Health Laboratory Fellowship Program: an APHL-CDC Initiative. The fellowship assignment is supported by Cooperative Agreement Number NU60OE000104 (CFDA No. 93.322), funded by the Centers for Disease Control and Prevention. This publication’s contents are solely the responsibility of the authors and do not necessarily represent the official views of the Centers for Disease Control and Prevention, the Department of Health and Human Services, or the Association of Public Health Laboratories and member laboratories. The fellowship assignment is 100% funded with federal funds from a federal program of $120,402,978.

---

**Supplementary Figure 1.**
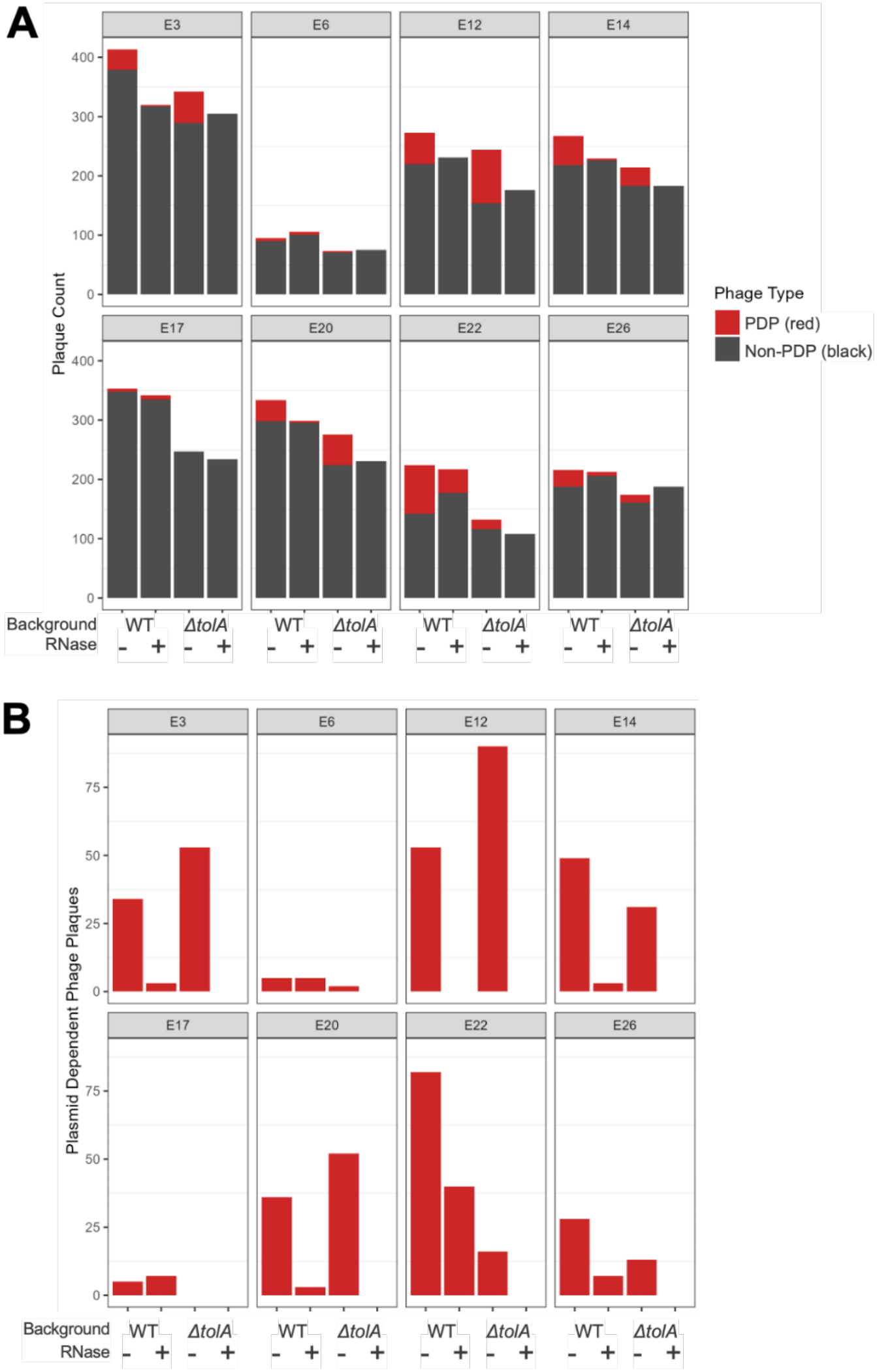
Recovery of plasmid-dependent and somatic phages from wastewater samples under different selective screening conditions. Plaques were enumerated from wastewater samples (designated E#) using F plasmid-dependent DisCo assays under four conditions: wild-type, RNase treatment, Δ*tolA*, and Δ*tolA* with RNase treatment. RNase treatment was used to inhibit ssRNA phages, whereas deletion of *tolA* prevented infection by canonical F-dependent inoviruses, enabling comparison of the relative contributions of these phage groups across samples. **(A)** Total plaque counts under each screening condition, including plasmid-dependent phages (red) and plasmid-independent phages (black). **(B)** Counts of plasmid-dependent phages only, highlighting variation in the relative abundance of RNase-sensitive and TolA-dependent phages between environmental samples.

**Supplementary Figure 2.**
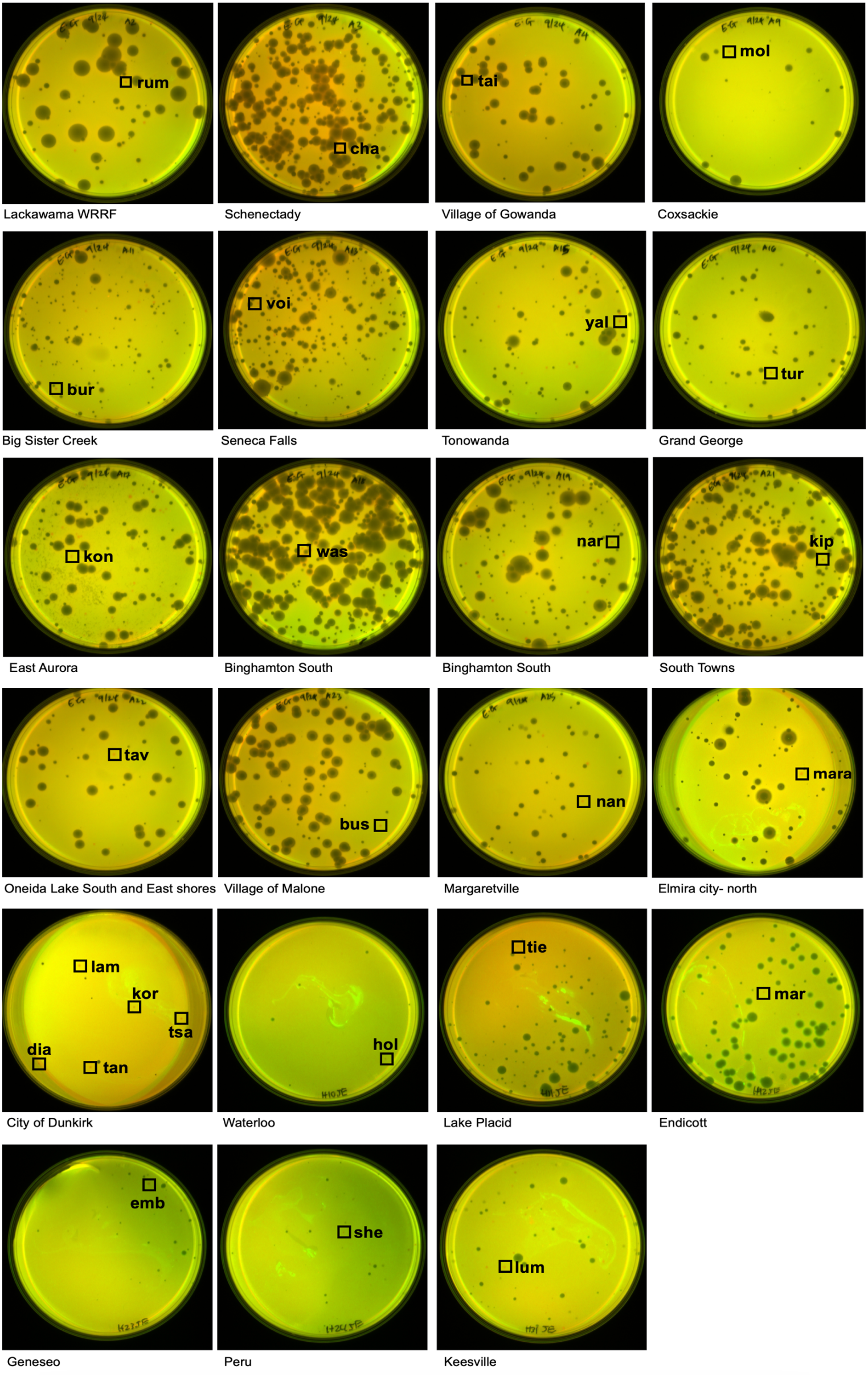
Original DisCo isolation plates for plasmid-dependent phages recovered in this study. Composite of the original DisCo plaque assays from which each phage in the collection was isolated. The plaque selected for subsequent purification is indicated by its corresponding phage name, with the wastewater sample of origin indicated beneath each image.

**Supplementary Figure 3.**
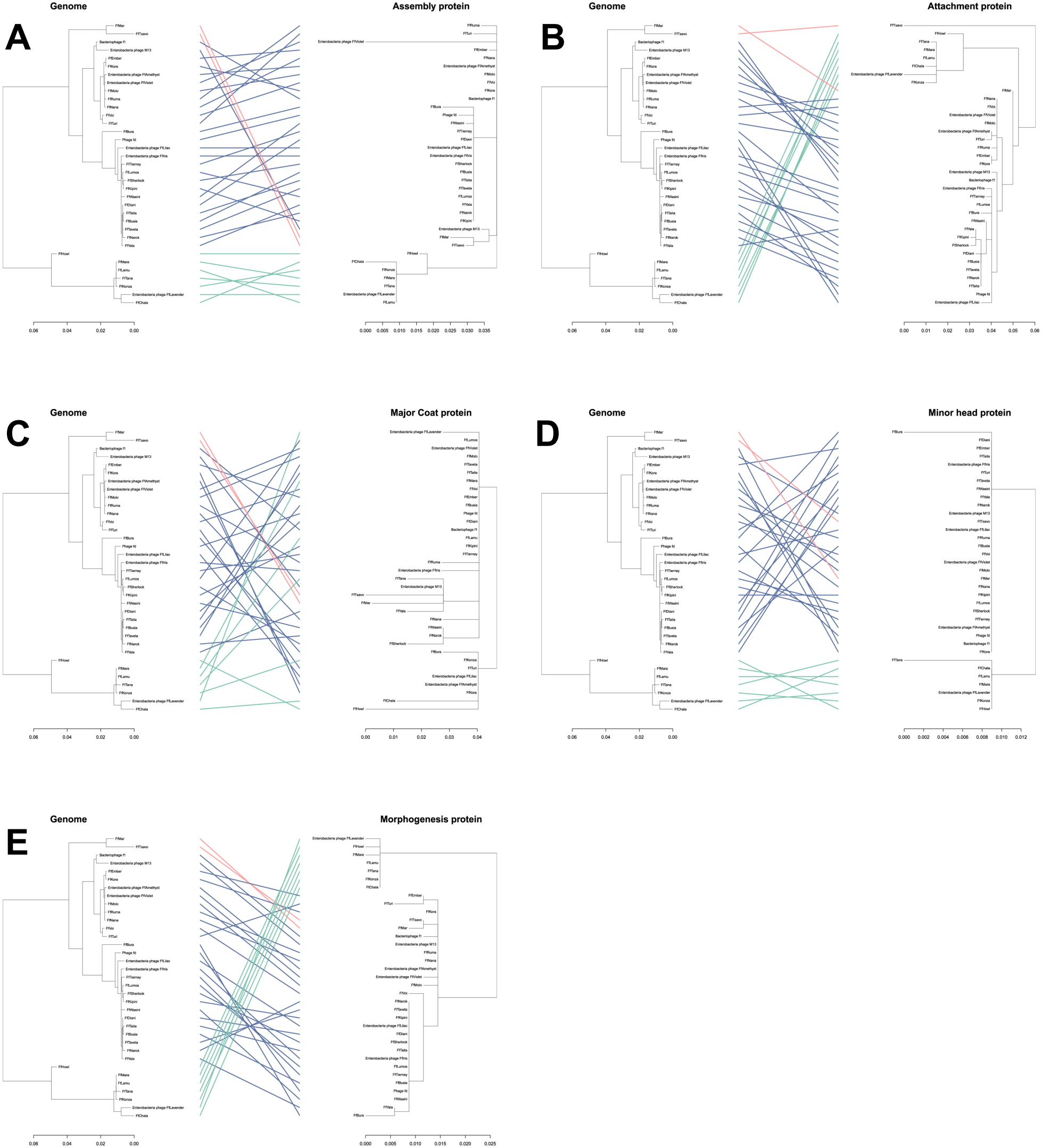
Comparison of whole-genome and individual gene phylogenies of plasmid-dependent inoviruses. Tanglegrams comparing phylogenetic relationships inferred from complete inovirus genomes with those inferred from five individual phage proteins: **(A)** assembly protein, **(B)** attachment protein, **(C)** major coat protein, **(D)** minor coat protein, and **(E)** morphogenesis protein. In each panel, the whole-genome phylogeny is shown on the left and the corresponding protein phylogeny on the right. Lines connect matching phages between trees and are colored according to the three major clades identified in the whole-genome phylogeny. Crossing lines indicate differences in phylogenetic topology between the whole-genome and individual protein trees.

