## Supplementary Tables for "Systematic discovery of plasmid-dependent inoviruses highlights broad infectivity across divergent IncF plasmids"

Table 1: Phages used in study

| Phage name | Short name | Lab code | Genome size | Isolation plasmid | Wastewater sample origin | Accession number |
|---|---|---|---|---|---|---|
| FfMara | mara | MP01 | 6412 | F | Elmira city- north | pending |
| FfTana | tan | MP02 | 6410 | F | City of Dunkirk | pending |
| FfLamu | lam | MP03 | 6412 | F | City of Dunkirk | pending |
| FfTsavo | sav | MP04 | 6393 | F | City of Dunkirk | pending |
| FfDiani | dia | MP05 | 6408 | F | City of Dunkirk | pending |
| FfKora | kor | MP06 | 6407 | F | City of Dunkirk | pending |
| FfRuma | rum | MP07 | 6406 | F | Lackawama WRRF | pending |
| FfChala | cha | MP08 | 6412 | F | Scenectady | pending |
| FfTaita | tai | MP09 | 6408 | F | Village of Gowanda | pending |
| FfMolo | mol | MP10 | 6407 | F | Coxsackie | pending |
| FfBura | bur | MP11 | 6408 | F | Big Sister Creek | pending |
| FfVoi | voi | MP12 | 6406 | F | Seneca Falls | pending |
| FfYala | yal | MP13 | 6408 | F | Tonowanda | pending |
| FfTuri | tur | MP14 | 6406 | F | Grand George | pending |
| FfKonza | kon | MP15 | 6412 | F | East Aurora | pending |
| FfWasini | was | MP16 | 6408 | F | Binghamton South | pending |
| FfNarok | nar | MP17 | 6408 | F | Binghamton South | pending |
| FfKipini | kip | MP18 | 6408 | F | South Towns | pending |
| FfTaveta | tav | MP19 | 6408 | F | Oneida Lake South and East shores | pending |
| FfBusia | bus | MP20 | 6408 | F | Village of Malone | pending |
| FfNana | nan | MP21 | 6407 | F | Margaretville | pending |
| FfHowl | how | H10 | 6408 | pED208 | Waterloo | pending |
| FfTierney | tie | H11 | 6409 | pED208 | Lake Placid | pending |
| FfMar | mar | H12 | 6405 | pED208 | Endicott | pending |
| FfEmber | emb | H23 | 6407 | pED208 | Geneseo | pending |
| FfSherlock | she | H24 | 6408 | pED208 | Peru | pending |
| FfLumos | lum | H31 | 6408 | pED208 | Keesville | pending |
| M13 | - | - | 6407 | F | - | NC_003287.2 |
| f1 | - | - | 6407 | F | - | NC_025823.1 |
| fd | - | - | 6408 | F | - | NC_025824.1 |
| FfAmethyst | ame | - | 6407 | F | - | PP430147.1 |
| FfIris | iri | - | 6408 | F | - | PP430148.1 |
| FfLavender | lav | - | 6412 | F | - | PP430149.1 |
| FfLilac | lil | - | 6424 | F | - | PP430150.1 |
| FfViolet | vio | - | 6388 | F | - | PP430151.1 |

Table 2: Wastewater sources

| Site description | Source type | Collection date | Phage isolated |
| --- | --- | --- | --- |
| Elmira city- north | Wastewater | 7/2/25 | FfMara |
| City of Dunkirk | Wastewater | 7/2/25 | FfTana |
| City of Dunkirk | Wastewater | 7/2/25 | FfLamu |
| City of Dunkirk | Wastewater | 7/2/25 | FfTsavo |
| City of Dunkirk | Wastewater | 7/2/25 | FfDiani |
| City of Dunkirk | Wastewater | 7/2/25 | FfKora |
| Lackawama WRRF | Wastewater | 9/10/25 - 9/11/25 | FfRuma |
| Scenectady | Wastewater | 9/11/25 | FfChala |
| Village of Gowanda | Wastewater | 9/11/25 | FfTaita |
| Coxsackie | Wastewater | 9/11/25 | FfMolo |
| Big Sister Creek | Wastewater | 9/11/25 | FfBura |
| Seneca Falls | Wastewater | 9/10/25 - 9/11/25 | FfVoi |
| Tonowanda | Wastewater | 9/11/25 | FfYala |
| Grand George | Wastewater | 9/11/25 | FfTuri |
| East Aurora | Wastewater | 9/11/25 | FfKonza |
| Binghamton South | Wastewater | 9/11/25 | FfWasini |
| Binghamton South | Wastewater | 9/11/25 | FfNarok |
| South Towns | Wastewater | 9/10/25 - 9/11/25 | FfKipini |
| Oneida Lake South and East shores | Wastewater | 9/11/25 | FfTaveta |
| Village of Malone | Wastewater | 9/11/25 | FfBusia |
| Margaretville | Wastewater | 9/11/25 | FfNana |
| Waterloo | Wastewater | 2/24/26 | FfHowl |
| Lake Placid | Wastewater | 2/24/26 | FfTierney |
| Endicott | Wastewater | | FfMar |
| Geneseo | Wastewater | 2/24/26 | FfEmber |
| Peru | Wastewater | 2/24/26 | FfSherlock |
| Keesville | Wastewater | 2/24/26 | FfLumos |

Table 3: Bacterial strains used in study

| Location | Strain background | Plasmids | Source | Description / construction |
|---|---|---|---|---|
| Owen lab 8.622 | E. coli MG1655 ΔtolA + F912 + pGreen (cmR) | pEB2-cmR-sGFP, F912 (kmR) | This study | MG1655 with *tolA* replaced by recombineering (pSIM5-tet) using cmR cassette from pKD3. cmR cassette flipped out using pCP20-tet. pEB2-cmR-sGFP transformed in via electroporation, then F912 plasmid from Quinones et al., 2024 delivered via conjugation. |
| Owen lab 8.623 | E. coli MG1655 ΔtolAc+ pRed (cmR) | pEB2-cmR-mScarlet-I | This study | MG1655 with *tolA* replaced by recombineering (pSIM5-tet) using cmR cassette from pKD3. cmR cassette flipped out using pCP20-tet. pEB2-cmR-mScarlet-I transformed in via electroporation. |
| Owen lab 8.621 | E. coli MG1655 F912 pGreen (cmR) | pEB2-cmR-sGFP, F912 | This study | MG1655 with pEB2-cmR-sGFP and F912 |
| Owen lab 8.620 | E. coli MG1655 pRed (cmR) | pEB2-cmR-mScarlet-I | This study | MG1655 with pEB2-cmR-mScarlet-I |
| Owen lab 8.624 | E. coli MG1655 ΔtolA + pGreen (cmR) pED208 | pEB2-cmR-sGFP, pED208 | This study | MG1655 Δ*tolA* pEB2-cmR-sGFP conjugated with pED208 plasmid |
| Owen lab 7.499 | E. coli BW25113 pEB2-cmR-sGFP + F912 | pEB2-cmR-sGFP, F912 (kmR) | This study | BW25113 pEB2-cmR-sGFP conjugated with F912 plasmid |
| Owen lab 7.491 | E.coli BW25113 pEB2-cmR-mScarlet | pEB2-cmR-mScarlet-I | Baym lab | |
| Owen lab 7.535 | E.coli W1485 ΔtolA::cmR | | This study | *tolA* replaced by recombineering (pSIM5-tet) using the cmR casette from pKD3 |
| Owen lab 6.441 | E. coli W1485 (F912) | F912 [finO::Plac-KanR-gfp.v912] | Baym Lab | |
| Owen lab 1.079 | E. coli MG1655 | | Hinton lab, Liverpool, UK | |
| Owen lab 6.484 | E. coli JE2571 pED208 | pED208 | Costa Lab, Imperial College London | |
| Owen lab 8.606 | E. coli BW25113 pEB2-cmR-sGFP + pED208 | pED208, pEB2-cmR-sGFP | This study | pED208 from E. coli JE2571 pED208 conjugated into E. coli BW25113 pEB2-cmR-sGFP |
| Owen lab 7.491 | E. coli BW25113 pEB2-cmR-mScarlet-I | pEB2-cmR-mScarlet-I | Baym Lab | |

Table 4: Plasmids used in study

| Plasmid name | Description | Reference |
|---|---|---|
| pKD3 | Template plasmid for amplifying cmR cassette for lambda red recombineering | Datsenko & Wanner, 2000 |
| pSIM5-tet | Temperature sensitive plasmid that encodes lambda red genes for recombineering | Datta et al., 2006 |
| pCP20-tet | Plasmid that encodes *flp* gene, a flippase that recombines the FRT sites in the recombineered FRT-antibioticR-FRT cassette, allowing the resistance cassette to be “flipped” out, leaving only an FRT scar. | Datsenko & Wanner, 2000 |

Table 5: Primers used in study

| Primer name | Sequence |
|---|---|
| del_tolA::cat_F | cttgcttgaaagagagcgggtaacaggcgaacagtttttggaaaccgagaCATATGAATATCCTCCTTAG |
| del_tolA::cat_R | ccgtggcaaccggtgcctgatgttgaccgtccgaacagtcaacatcgcgaGTGTAGGCTGGAGCTGCTT |
| tolA_ext_F | cgcagcctatctaaacatct |
| tolA_ext_R | ccgatggcccacgataattt |
